# The Integrated Stress Response Pathway Improves Aged Murine Muscle Stem Cell Activation and *in vivo* Regeneration

**DOI:** 10.1101/2025.09.26.678822

**Authors:** Alexander D. Brown, Annarita Scaramozza, Hanzhi Zhang, Samiha Mahin, Nivedita Suresh, Takeshi Tsusaka, Susan Eliazer, Xuhui Liu, Brian Feeley, Andrew S. Brack

## Abstract

For efficient regeneration, muscle stem cells (MuSCs) transition out of quiescence through a series of progressively more activated states. During MuSC aging, transition through the earliest steps is the slowest and delayed, with the molecular regulators that govern this transition not well characterized. By analyzing the dynamic changes of MuSCs at the molecular (scRNA-Seq and Cell Painting) and phenotypic (heteromotility) level at single cell resolution we found that the Integrated Stress Response (ISR) Pathway is a critical regulator of MuSC transition states. Aged MuSCs have increased baseline ISR activity in quiescence that does not increase during activation to levels observed in adult MuSCs. Rapid and transient pharmacological ISR activation *in vitro* was sufficient to increase aged MuSC activation rate and migratory behavior as well as alter the transcriptional states toward a younger phenotype. ISR activation also improved aged MuSC potency and aged mouse muscle regeneration *in vivo*. Therefore, pharmacological activation of the ISR has therapeutic potential to improve MuSC function and skeletal muscle repair during aging.

## INTRODUCTION

Adult stem cells are key regulators of tissue regeneration after injury. Skeletal muscle regeneration is dependent on muscle stem cells (MuSCs), also known as satellite cells (Lepper *et al*, 2011; Murphy *et al*, 2011; Sambasivan *et al*, 2011). Upon injury, MuSCs exit a quiescent state, activate, and expand to produce progenitor cells that rebuild muscle fibers (Brack & Rando, 2012). The transition from quiescence to activation is characterized by a series of molecular, morphological, and phenotypic events, which can be defined as transition states (Kimmel *et al*, 2020; Raval *et al*, 2020; Ryall *et al*, 2015; Tang & Rando, 2014). Single-cell RNA (scRNA) sequencing has revealed transcriptional heterogeneity across the MuSC compartments in murine and human muscle (Barruet *et al*, 2020; De Micheli *et al*, 2020; Dell’Orso *et al*, 2019; Kimmel *et al*, 2020; Raval *et al*, 2020; Rubenstein *et al*, 2020). Trajectory analysis of single-cell transcriptomes from adult murine MuSCs revealed two sequential states formed in response to activation cues *in vitro*, one in a more activated state than the other (Kimmel *et al*, 2020). RNA velocity revealed a non-linear change in transcription across the activation space, consistent with a rate-limiting step (Kimmel *et al*, 2020). Analysis of the cell cycle and motility at the single-cell level confirm that the earliest state transitions are the slowest (Kimmel *et al*, 2020; Raval *et al*, 2020).

Extrinsic stimuli that liberate MuSCs from quiescence into a more primed state can accelerate muscle regeneration (Eliazer *et al*, 2019; Rodgers *et al*, 2014, 2017; Schröder & Kaufman, 2005). We have previously shown that pre-defined MuSC subsets with high regenerative potential (higher potency), such as label retaining MuSCs (LRCs) or Pax3^+^ MuSCs traverse through activation states more rapidly (Chakkalakal *et al*, 2012; Kimmel *et al*, 2020; Scaramozza *et al*, 2019). These data suggest that stem cell potency is related to the rate at which MuSCs transition through activation states.

In aged muscle, MuSC function is impaired resulting in delayed muscle regeneration (Bernet *et al*, 2014; Brack *et al*, 2007; Conboy *et al*, 2003; Cosgrove *et al*, 2014; García-Prat *et al*, 2016; Sousa-Victor *et al*, 2014). Aged MuSCs activate slowly and are susceptible to apoptosis, suggesting an inappropriate response to activation cues and loss of resilience (Liu *et al*, 2018). Cell cycle entry kinetics, cell behavior, and transcriptional analyses reveal that in response to mitogenic signals, aged MuSCs traverse activation states more slowly than adult MuSCs (Kimmel *et al*, 2020). While these data suggest that the loss in potency of aged MuSCs is related to aberrant cell state transitions, the molecular regulators driving this process are not yet defined.

In this study, we generated multimodal single cell data and leveraged publicly available single cell MuSC transcriptomic data to model state transitions to investigate the mechanism driving MuSC activation dynamics from adult and aged murine muscle (Kimmel *et al*, 2020). We found that the integrated stress response (ISR) pathway was dynamically expressed during activation of adult MuSCs. Using pharmacological approaches to modulate this pathway, we show that the ISR pathway is required for adult MuSC activation. Transient ISR activation accelerates aged MuSC activation, promotes survival *in vitro*, and enhances stem cell potency and muscle repair *in vivo*. Therefore, modulation of ISR pathway represents a promising therapeutic approach to enhance MuSC function and increase cellular resilience during aging.

## RESULTS

### The ISR pathway undergoes dynamic expression changes during adult MuSC activation

To identify dynamically altered gene expression patterns governing MuSC activation, we reanalyzed our scRNA-sequencing data from freshly isolated MuSCs (quiescence (Q)) and MuSCs after 18 hours of *in vitro* activation (A), within which supervised clustering and UMAP projection of transcriptional state space revealed four distinct transcriptional clusters (Figure 1A, B). Pseudotime lineage reconstruction suggested cells followed a series of sequential states from cluster 1 to cluster 4, with cells in cluster 4 representing the most activated state (Kimmel *et al*, 2020).

**Figure 1.**
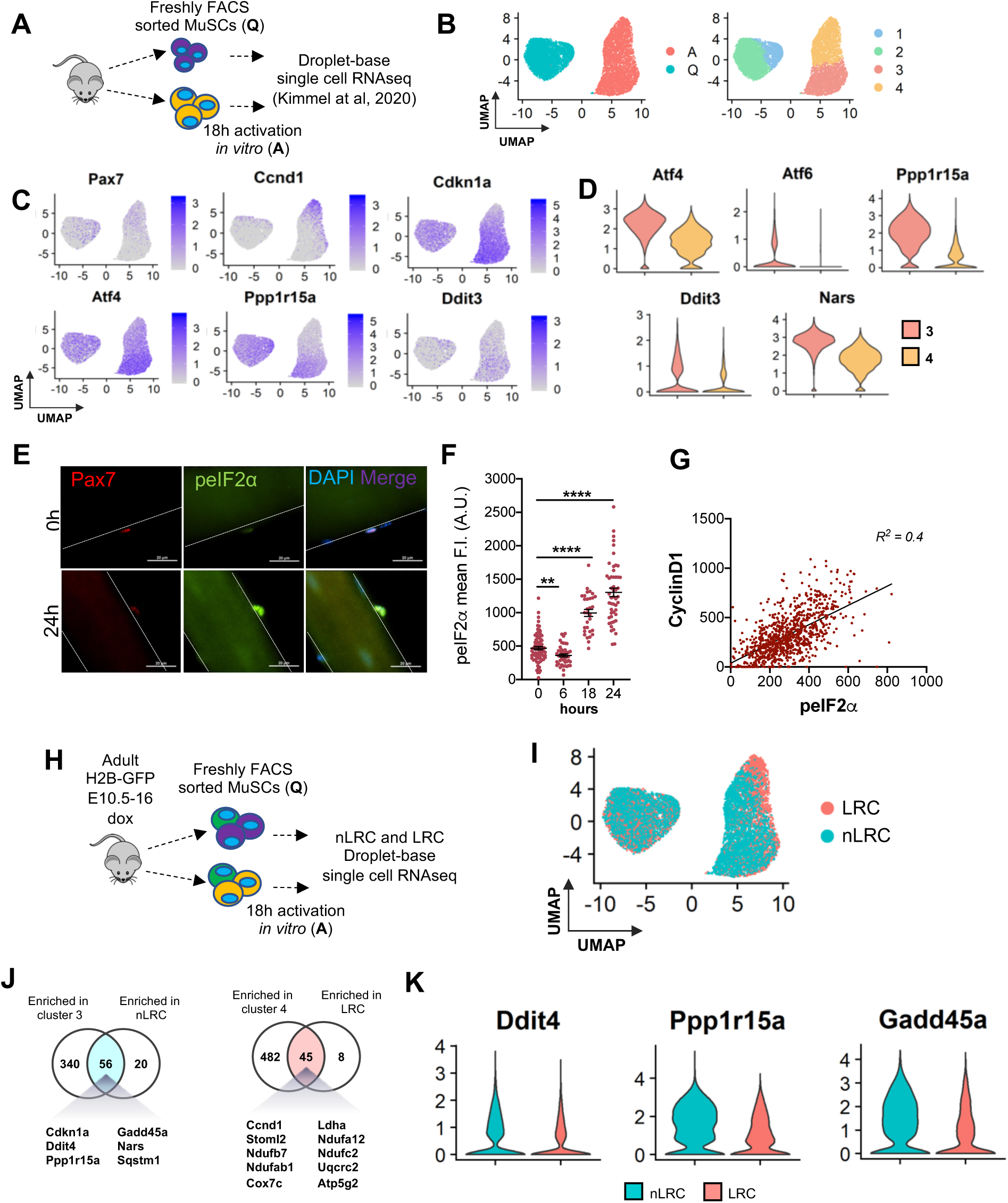
The ISR pathway undergoes dynamic expression changes during adult MuSC activation. **A.** Schematic of single-cell RNA sequencing of freshly FACS sorted (Q=quiescence) and 18 hours *in vitro* activated (A=activation) from adult mouse adapted from Kimmel et al., 2020. n=2 mice. **B.** UMAP projection of transcriptional state space (left) and unsupervised clustering showing two transcriptional clusters within the Q-MuSC and A-MuSC populations (right). **C.** Overlay of expression of cell cycle genes and ISR-related genes in scaled log (UMI+1) on the UMAP plot. **D.** Violin plots of ISR-related genes significantly different among cluster 3 and cluster 4. **E.** Representative images of single muscle fibers from adult mice (3-6 months) stained for Pax7 (red), p-eIF2α (green) and DAPI (blue) immediately after isolation (0 hours) and 24 hours later. Scale bar, 20 µm. **F.** Time course of p-eIF2α mean fluorescence intensities in Pax7^+^ MuSCs on adult single muscle fibers at 0, 6, 18 and 24 hours after isolation. Mean±SEM, n=3 mice, **p<0.01, ****p<0.001. **G.** Correlation between CyclinD1 and p-eIF2α (R^2^=0.4) after 18 hours *in vitro.* n=3 mice. **H.** Schematic of single-cell RNA sequencing of FACS sorted (quiescent and activated) label retaining cells (LRCs) and non label retaining cells (nLRCs) from TetO-H2B-GFP transgenic mouse model. n=2 mice. **I.** UMAP projection of unsupervised clustering showing nLRC and LRC transcriptional clusters within the quiescent and activated adult MuSC populations. **J.** Venn diagrams showing common genes with statistically significant differentially expressed genes among cluster 3 and nLRC (left) and between cluster 4 and LRC (right) (13% and 8% transcriptional overlap). **K.** Violin plots of ISR-related genes differentially expressed among nLRC and LRC.

To identify the genes dynamically regulated during activation, we analyzed differentially expressed genes between cluster 3 and cluster 4 (Figure 1C). Cluster 3 had more *Cdkn1a* and Cluster 4 had more *Pax7* and *Ccnd1*. Gene ontology analysis revealed genes associated with RNA processing, tRNA processing, and ribonucleoprotein biogenesis were enriched in cluster 3 MuSCs, while metabolites, oxidation-reduction, and ATP metabolic, and mitochondrion organization processes were enriched in cluster 4 MuSCs (Figure S1A). Therefore, cluster 4 represents MuSCs that are in a more advanced activation state.

Pathway analysis indicated genes associated with the ISR pathway were upregulated during the transition from quiescence to activation (Figure 1C; S1B). The ISR pathway is an adaptive pathway activated in response to diverse cellular stressors to restore cellular homeostasis (Costa-Mattioli & Walter, 2020; Pakos-Zebrucka *et al*, 2016) and has been implicated in adult MuSC self-renewal (Zismanov *et al*, 2016). Analysis of the ISR pathway revealed that *Activated transcription factor 4 (ATF4)*, *DNA damage inducible transcript 3 (Ddit3)* and *growth arrest and DNA damage-inducible protein* (*GADD34* also known as *Ppp1r15a*) expression was higher in cluster 3 (Figure 1D; S1C-D). Therefore, downstream activators and feedback inhibitors of the ISR pathway are expressed in cluster 3, which represents MuSCs that are in a less activated state.

To activate the ISR, stress signals trigger reversible phosphorylation of the alpha subunit of eukaryotic translation initiation factor 2 (eIF2α) on serine 51, which halts translation and global protein synthesis, while allowing selective translation of genes involved in cell survival and recovery (Ron, 2002). To determine the kinetics of ISR signaling activity in adult MuSCs, we performed a time course analysis of p-eIF2α expression on single muscle fibers during activation, which occurs progressively upon isolation. Freshly isolated quiescent MuSCs express moderate p-eIF2α, which decreases at 6 hours (Zismanov *et al*, 2016) and progressively increased at 18 and 24 hours after isolation (Figure 1E-F). At 24 hours, while p-eIF2α was significantly higher on average, the greater heterogeneity in expression suggests that the ISR could be dampened in some MuSCs compared to others (Figure 1F). To compare ISR signaling within the activated clusters, we stained MuSCs after 24 hours *in vitro* for p-eIF2α and CyclinD1, which we showed was highest in cluster 4 (Figure 1C). We found a positive correlation between p-eIF2α and Cyclin D1 staining (R^2^ = 0.4), demonstrating that activation of the ISR corresponded with Cyclin D1 expression (Figure 1G).

To compare ISR signaling between adult MuSCs with different activation rates and potencies, we used the TetO-H2B-GFP transgenic mouse model, which serves as a readout of proliferative kinetics (Figure 1H) (Chakkalakal *et al*, 2014; Kimmel *et al*, 2020). Similar to MuSC cluster 4, label-retaining MuSCs (LRCs) activate more rapidly than non-LRCs (nLRCs) and we found they share some transcriptional overlap (8%) with cluster 4 (Kimmel *et al*, 2020) (Figure 1I). Venn diagrams show that there were overlapping gene sets including Stress Response and Protein Misfolding in cluster 3 and nLRCs and ATP Metabolism and Oxidative Metabolism between cluster 4 and LRCs (Figure 1J). Importantly, we found multiple ISR genes enriched in activated nLRCs compared to LRCs (Figure 1K; S1E). Therefore, ISR genes are transiently upregulated in cluster 3 (predominately nLRCs), the less activated state and dissipate in cluster 4 (predominately LRCs), the more activated state.

### The ISR is associated with activation of adult MuSCs

Because ISR genes are enriched in the less activated MuSCs, we speculated that either the ISR inhibits state transitions, maintaining cells in a less advanced activation state, or the ISR is required for early state transitions and dissipates when MuSCs transition to more advanced activation state. To test these possibilities, we stained fluorescence activated cell sorted (FACS) MuSCs with markers of activation following exposure to an ISR activator, Sal003 (ISR-a) (Costa-Mattioli & Walter, 2020) or an ISR inhibitor ISRIB (ISR-i) (Sidrauski *et al*, 2013).

Using EdU incorporation as a readout of MuSC activation, adult muscle fibers treated with ISR-a had a higher percentage of EdU^+^ MuSCs, whereas ISR-i treated MuSCs had a lower fraction of EdU^+^ cells (Figure 2A). After 48 hours in ISR-a, the fraction of apoptotic (Casp3^+^) MuSCs did not change, while ISR-i treatment increased the fraction of apoptotic MuSCs (Figure 2B), suggesting that mounting an ISR response during activation is critical for MuSC survival.

**Figure 2.**
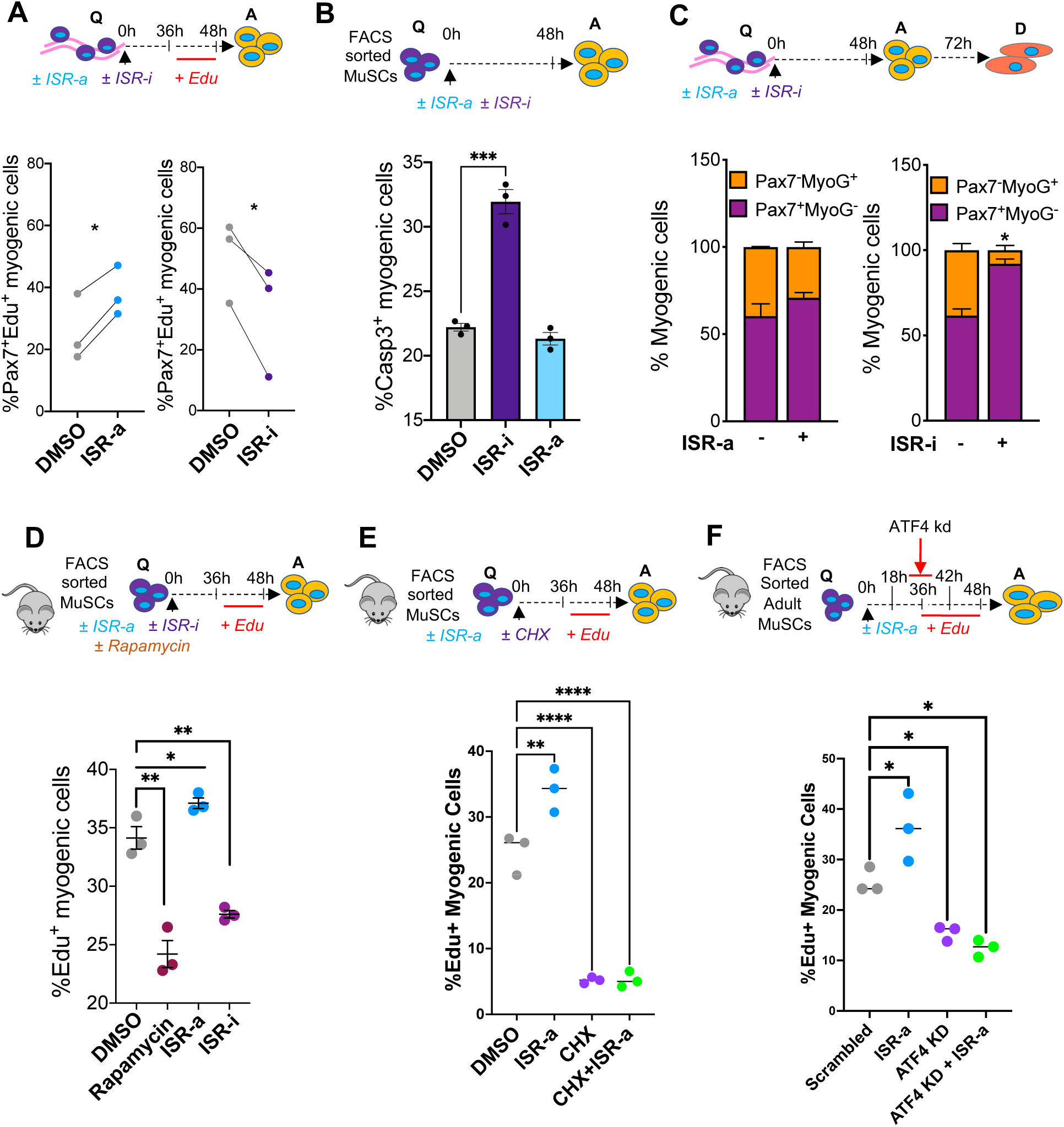
Stimulating the ISR pathway is associated with accelerated adult MuSC activation through selective translation. **A.** Schematic of single muscle fibers from adult mice (3-6 months) treated either with DMSO, 10 µM Sal003 (ISR-a) or 200 nM ISRIB (ISR-i), pulsed with EdU and analyzed at 48 hours in culture (top); Q = quiescent, A = activated. Percentage of Pax7^+^EdU^+^ cells at 48 hours after ISR-a (bottom left) and ISR-i treatment (bottom right). Mean±SEM; n=3; *p<0.05. **B.** Schematic of FACS sorted adult MuSCs *in vitro* cultured in DMSO, ISR-a or ISR-i for 48 hours (top). Percentage of Casp3^+^ myogenic cells at 48 hours after treatment (bottom). Mean±SEM; n=3; ***p<0.001. **C.** Schematic of single muscle fibers from adult mice (3-6 months) treated either with DMSO, ISR-a or ISR-i and cultured for 72 hours in culture (top); Q = quiescent, A = activated, D = differentiated. Percentage of Pax7^-^/MyoG^+^ (orange) and Pax7^+^/MyoG^-^ (purple) cells at 72 hours in culture after ISR-a treatment (bottom left) and ISR-i treatment (bottom right). Mean±SEM; n=2, *p<0.05. **D.** Schematic of FACS sorted adult MuSCs *in vitro* cultured in DMSO, 10 µM rapamycin, ISR-a or ISR-i for 48 hours (top); Q = quiescent, A = activated. Percentage of EdU^+^ FACS sorted myogenic cells at 48 hours after DMSO, Rapamycin, ISR-a or ISR-i treatment. Mean±SEM; n=3; *p<0.05; **p<0.01. **E.** Schematic of FACS sorted adult MuSCs *in vitro* cultured in DMSO, ISR-a, 100 µg/ml cycloheximide (CHX), or CHX+ISR-a for 48 hours (top); Q = quiescent, A = activated. Percentage of EdU^+^ FACS sorted myogenic cells at 48 hours after DMSO, ISR-a, CHX, or CHX+ISR-a treatment. Mean±SEM; n=3; *p<0.05; **p<0.01, ***p<0.001. **F.** Schematic of FACS sorted adult MuSCs *in vitro* cultured in ISR-a pulsed with EdU and ATF4 shRNA for 48 hours (top). Percentage of Pax7^+^EdU^+^ cells at 48 hours after ISR-a, ATF4 shRNA or with ISR-a+ATF4 shRNA (bottom). Mean±SEM; n=3; *p<0.05.

At 72 hours, the number of myogenic cells per muscle fiber increased after ISR-a and decreased after ISR-i treatment, coinciding with increased apoptosis (Figure 2B; S2A), suggesting that stimulating the ISR accelerated MuSC activation and proliferation without affecting survival or commitment (Figure 2B-C). After 72 hours of ISR-i treatment, a greater fraction of MuSCs were Pax7^+^ compared to control, consistent with limited activation and lineage progression (Figure 2C). Together these results suggest that ISR stimulation is required for adult MuSC activation whereas ISR inhibition delays activation and promotes apoptosis.

### The ISR accelerates MuSC activation through selective translation

Activation of the ISR halts global protein synthesis, while allowing for selective mRNA translation. Global protein translation promotes the transition of quiescent MuSCs toward a primed state (Rodgers *et al*, 2014). To determine whether the effect of ISR on MuSC activation can be ascribed to selective translation or global protein synthesis inhibition, we determined whether there was any cross regulation between ISR and mTORC1 activity. FACS sorted MuSCs from adult mice were treated with the mTOR inhibitor rapamycin, ISR-a, or ISR-i for 3 hours. Rapamycin reduced pS6 with no effect on p-eIF2α, and ISR-a treatment increased p-eIF2α without any effect on pS6. In contrast, ISR-i increased pS6, suggesting a possible interaction between ISR and mTORC1 signaling (Figure S2B). Next, we determined whether rapamycin affected MuSC activation. In contrast to ISR-a, rapamycin decreased the percentage of EdU^+^ MuSCs (Figure 2D). We also treated MuSCs with cycloheximide (CHX), another inhibitor of global protein synthesis for 3 hours, followed by a 45 hour wash out in growth media. CHX decreased the percentage of EdU^+^ MuSCs compared to controls, suggesting MuSC activation is impaired when global synthesis is inhibited. We also found that ISR-a treatment did not rescue the CHX-mediated decline in MuSC activation (Figure 2E), suggesting that ISR is not sufficient to activate MuSCs in the absence of global protein synthesis.

Finally, we asked whether the effect of ISR on MuSC activation was through ATF4. *Atf4* knockdown in MuSCs decreased the percentage of EdU^+^ MuSCs and ISR-a treated MuSCs increased the percentage EdU^+^ cells which was abrogated after *Atf4* knockdown (Figure 2F). ISR-a did not rescue the *Atf4*-mediated decline in MuSC activation, consistent with *Atf4* acting downstream of p-eIF2α (Figure 2F). Together, these data suggest that induction of the ISR accelerates MuSC activation through selective translation and is dependent on *Atf4*.

### Activation of the ISR is sufficient to accelerate aged MuSC activation

Adult and aged MuSCs transition through the same activation states, but aged MuSCs do so more slowly, spending more time in a less activated state (Kimmel *et al*, 2020). Like adult activated MuSCs, we observed an enrichment of ISR pathway genes in the less activated MuSC state from aged muscle (Figure S3A). Aged quiescent MuSCs had elevated baseline p-eIF2α compared to adult quiescent MuSCs, but the rapid increase in p-eIF2α expression during adult MuSC activation was abrogated in aged MuSCs (Figure 3A). This implies that aged MuSCs are not able to induce an ISR response during activation, unlike adult MuSCs.

**Figure 3.**
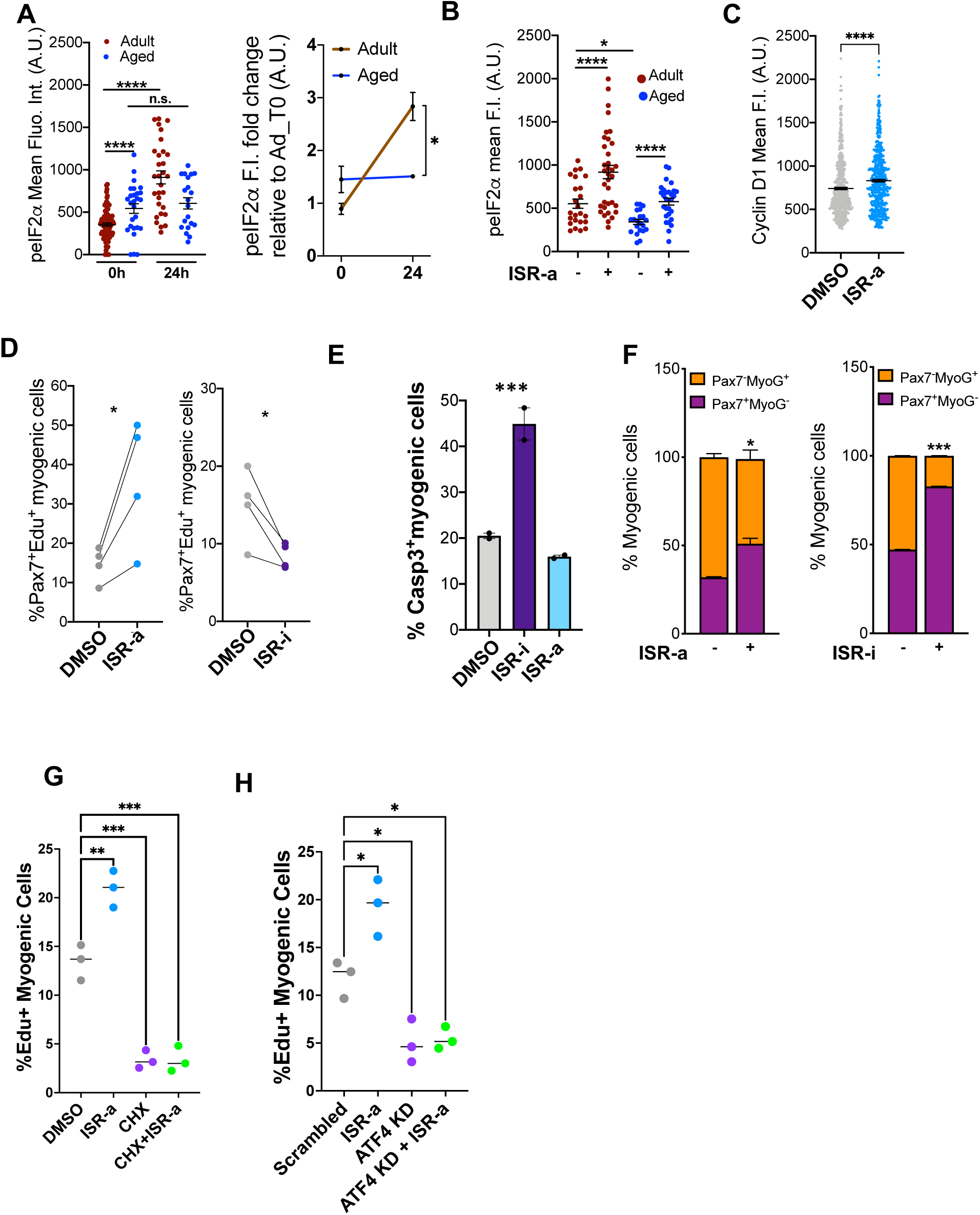
Activation of ISR is sufficient to restore function of aged MuSCs. **A:** Time course (left) and relative fold change to adult (right) of p-eIF2α mean fluorescence intensities in Pax7^+^ MuSCs on aged single muscle fibers at 0 and 24 hours after isolation. Mean ± SEM, n = 3 mice, *p<0.05, ****p<0.001, ns=nonsignificant. **B:** p-eIF2α mean fluorescence intensities in Pax7^+^ MuSCs on aged single muscle fibers with DMSO or ISR-a at 24 hours after isolation. Mean±SEM, n=3 mice, *p<0.05, ****p<0.001. **C:** CyclinD1 expression in aged MuSCs with DMSO or ISR-a after 18 hours *in vitro.* Mean±SEM; n=3; *p<0.05. **D:** Percentage of Pax7^+^EdU^+^ cells at 48 hours after ISR-a (left) and ISR-i treatment (right). Mean±SEM; n=3; *p<0.05. **E.** Schematic of FACS sorted aged MuSCs *in vitro* cultured in presence or absence of ISR-a and ISR-b for 48 hours. Percentage of Casp3^+^ myogenic cells at 48 hours after DMSO, ISR-a and ISR-i treatment (bottom). Mean±SEM; n=2; ***p<0.001. **F:** Percentage of Pax7^-^/MyoG^+^ (orange) and Pax7^+^/MyoG^-^ (purple) cells at 72 hours in culture after ISR-a treatment (left) and ISR-b treatment (right). Mean±SEM; n=2, *p<0.05, ***p<0.001 **G:** Percentage of EdU^+^ FACS sorted myogenic cells at 48 hours after DMSO, ISR-a, CHX, or CHX+ISR-a treatment. Mean±SEM; n=3; **p<0.01, ***p<0.001. **H:** Percentage of Pax7^+^EdU^+^ cells at 48 hours after ISR-a, ATF4 shRNA or with ISR-a+ATF4 shRNA. Mean±SEM; n=3; *p<0.05.

Because aged quiescent MuSCs have higher baseline p-eIF2α but remain mostly quiescent, we tested whether aged MuSCs are competent to respond to increased ISR activity. Single muscle fibers from adult and aged mice were cultured in DMSO or ISR-a for 24 hours. ISR-a restored p-eIF2α levels in aged activated MuSCs to that of adult levels (Figure 3B). CyclinD1 expression also increased in the presence of ISR-a, suggesting that forced ISR activation can drive aged MuSCs towards a more advanced activated state (Figure 3C).

Phenotypically, ISR-a treatment increased the fraction of activated and decreased the fraction of apoptotic aged MuSCs, whereas ISR-i had the opposite effect (Figure 3D-E). Cell fate analysis revealed that ISR-a increased the percentage of Pax7^+^ aged MuSCs at the expense of committed Myogenin^+^ cells compared to DMSO controls, suggesting MuSC expansion without differentiation (Fujita *et al*, 2021; Zismanov *et al*, 2016). Treatment with ISR-i retained aged MuSCs in a Pax7^+^/Myogenin^-^ fate, presumably due to stalled proliferation and cell death (Figure 3F). Global protein synthesis was blocked with CHX and led to a lower percent of EdU^+^ aged myogenic cells compared to control with no rescue when co-treated with ISR-a (Figure 3G). In aged MuSC activation, we asked whether the ISR pathway signaled in a canonical manner through *Atf4*. *Atf4* knockdown decreased the percent of EdU^+^ positive cells and ISR-a did not rescue this inhibition (Figure 3H). Taken together, inducing the ISR promoted aged MuSC activation through selective translation and in a *Atf4* dependent manner.

### ISR activation induced adult-like behavior in aged MuSC through alterations in actin and mitochondria morphology

Because activation was improved in aged MuSCs with ISR-a, we tested whether ISR-a altered aged MuSC migration behavior within the first 48 hours using our heteromotility assay (Figure 4A: (Kimmel *et al*, 2018)). We revealed that aged MuSCs with ISR-a improved cell linearity (ability of the cell to directly migrate from its start to end position) beyond the level of the adult (Figure 4B). Further, ISR-a partially improved features of cell movement (speed, distance, time moving, progressivity) and cell direction (theta, autocorrelation, hurst exponent, proportion of right turns) (Figure S4A: heteromotility definitions). We next determined whether ISR-a treatment induces aged MuSCs to transition to a more adult-like motile state. Clustering showed overlap between adult, aged, and aged treated with ISR-a (Figure 4C), with three distinct behavioral states (Figure 4D, E). Across all experimental groups, most cells were in cluster 2 during the 48 hours of activation (Figure 4D) and aged MuSCs had fewer cells in cluster 1. ISR-a aged MuSCs had a higher fraction of MuSCs in cluster 1, at the expense of aged MuSCs in cluster 3, trending to more adult-like behavior (Figure 4D). Therefore, ISR-a increased the proportion of aged MuSCs from cluster 3 to 1, where cells spent more time moving, with greater speed and distance (Figure 4E), an indication of youthful behavior *in vitro*.

**Figure 4.**
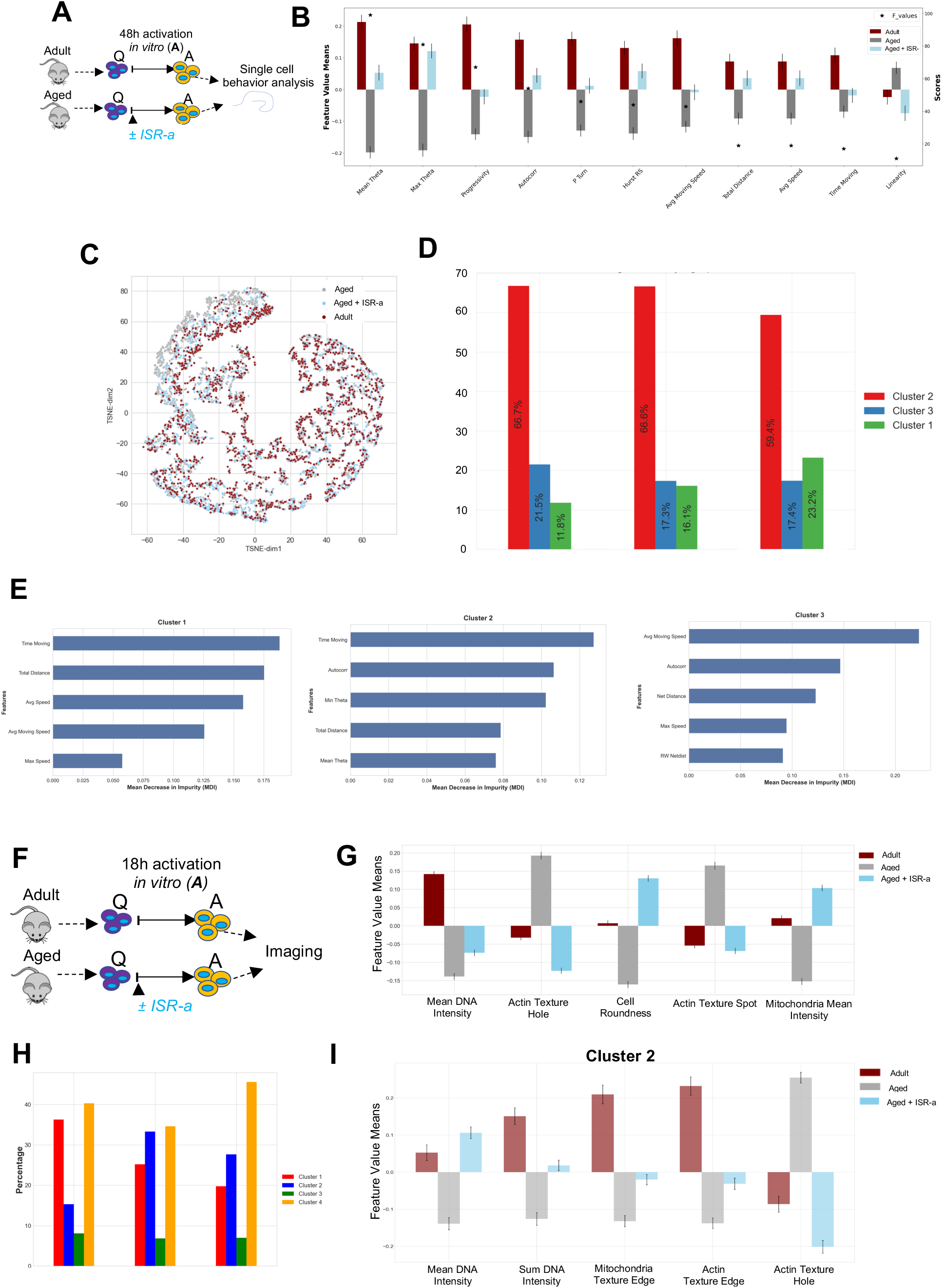
Aged MuSC behavior was improved with ISR activation through alterations in actin and mitochondria morphology. **A:** Schematic of FACS sorted adult and aged MuSCs *in vitro* cultured and live imaged in DMSO or ISR-a for 48 hours. Images were processed through our heteromotility pipeline. **B:** MuSC heteromotility features in adult DMSO, aged DMSO and aged ISR-a. Mean±SEM; n=3; *p<0.05. **C:** TSNE projection showing activated MuSC states in adult DMSO, aged DMSO and aged ISR-a. **D:** Percentage of adult DMSO, aged DMSO and aged ISR-a in three distinct MuSCs clusters. **E:** Heteromotility features ranked on importance within cluster 1 (left), 2 (middle) and 3 (right). **F.** Schematic of FACS sorted adult and aged MuSCs cultured in presence or absence of ISR-a for 18 hours. **G.** ANOVA F-Value test showing most variable feature means of all cell population between adult DMSO, aged DMSO, and aged ISR-a. n=3. **H.** Percentage of clusters within cell populations from PCA projection of morphological state space and unsupervised GMM clustering. **I.** Histograms of the most variable features means in cluster 2 between adult DMSO, aged DMSO and aged ISR-a.

Next, we leveraged the cell painting assay (Bray *et al*, 2016) to explore the organelle characteristics and morphology in ISR-a-treated aged MuSCs during activation (Figure 4F). In aged MuSCs, DNA and mitochondrial intensity, actin texture hole and spot (non-homogenous actin structure) were restored back to the level of adult MuSCs by ISR-a treatment (Figure 4G: S4D). Gaussian Mixture Models identified four distinct clusters which were visualized onto a principal component analysis (PCA) plot of adult, aged, and ISR-a-treated aged MuSCs with cluster 2 and 4 having the highest proportion of ISR-a aged MuSCs (Figure 4H, S4B). Within cluster 2, DNA staining intensity, mitochondria texture edge (mitochondrial structures), actin texture edge (actin structures along the cell perimeter) and actin texture hole were partially or fully restored back to untreated adult MuSCs (Figure 4I). Cluster 4 had greater DNA staining intensity, actin texture spot, and brightness (high actin intensity) in ISR-a-treated aged versus untreated aged MuSCs (Figure S4C). In this cluster, ISR-a treatment resulted in significantly greater mitochondrial staining intensity compared to both adult and aged. Our data therefore suggest that ISR-a in aged MuSCs drives behavior through changes in actin and mitochondrial dynamics towards a more adult-like state.

### ISR activation in aged MuSCs is molecularly restored to an adult state

We next asked whether transcriptional changes drive the structural alterations observed with ISR-a treatment. Adult and aged MuSCs were activated for 18 hours with either DMSO or ISR-a, re-sorted to collect live cells, and sequenced (Figure 5A). Comparing activated adult, aged, and aged with ISR-a treatment, UMAP projection identified three transcriptionally unique clusters (Figure 5B and S5A). Cluster 0 comprised a similar number of adult and aged ISR-a-treated cells while clusters 1 and 2 were mostly made up of adult cells. Pseudotime analysis showed that cluster 0 was less advanced compared to cluster 1 and 2 (Figure 5B). Cluster 0 had a higher expression of ISR genes, which was consistent with the less activated cluster in Figure 1C. Cluster 1 and 2 had a lower expression of ISR genes and higher expression of *Ccnd1*, consistent with the more activated cluster. Across experimental groups, ISR-a treatment increased the fraction of cells in the most activated cluster, cluster 2 (Figure 5C, D). Pairwise comparisons of the three transcriptionally unique clusters by GO analysis revealed that cluster 0 had consistently high expression of genes characteristic of a less activated state (Figure 5E). Cluster 1 exhibited high expression of genes related to mitochondrial translation and aerobic respiration aligning with a more activated state. However, while cluster 1 and 2 appeared more active, cluster 1 also had high expression of genes associated with less activated state compared to cluster 2, which showed high expression of genes associated with a more activated state (Figure S5B). Together, these data demonstrate that ISR-a treatment accelerates the transition of aged MuSCs through activation state space.

**Figure 5.**
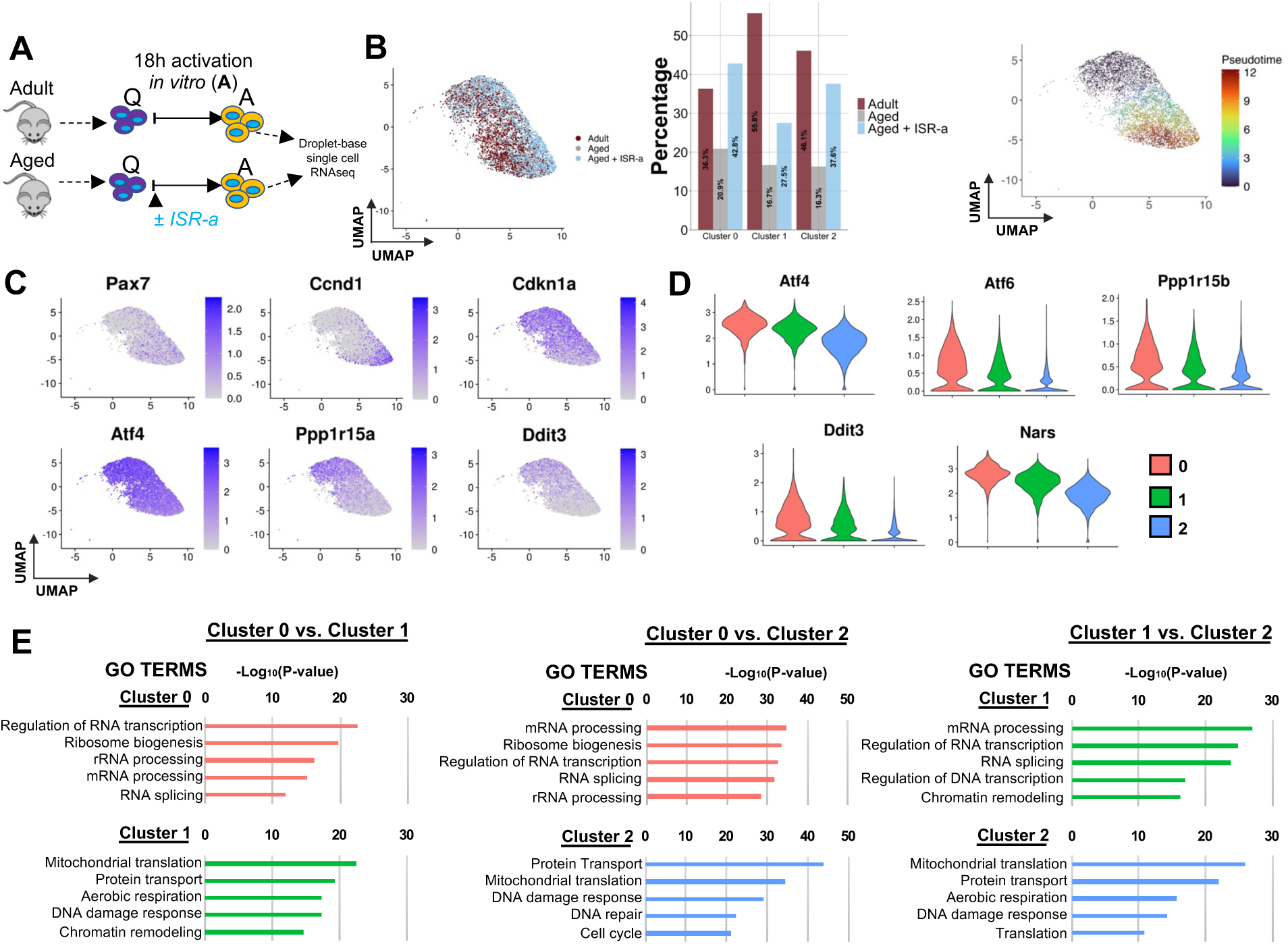
ISR activation in aged MuSCs molecularly and morphologically recapitulates adult MuSCs. **A.** Schematic of single-cell RNA sequencing of FACS sorted adult and aged MuSCs in the presence or absence of ISR-a and activated (A) for 18 hours *in vitro.* n=2 mice. **B.** UMAP projection of transcriptional state space (left) showing three transcriptional clusters within the activated adult, aged, and aged ISR-a cell populations. Percentage of cell populations (adult, aged, aged ISR-a) within clusters (middle). UMAP projection of psuedotime analysis of each cluster (right). **C.** Overlay of expression of cell cycle genes and ISR-related genes in scaled log (UMI + 1). **D.** Violin plots of ISR genes significantly different among cluster 1 compared to cluster 0 and cluster 2. **E.** Gene ontology analysis for common genes with statistically significant (p<0.05) differentially expressed genes among pairwise comparisons of all clusters were performed using DAVID, only considering biological processes.

### Activating the ISR in aged MuSCs improves transplant potential and regulates the more primed MuSC subset

Because MuSC transplantation potential decreases with aging, we tested whether ISR modulation could influence this effect (Bernet *et al*, 2014; Chakkalakal *et al*, 2012; Cosgrove *et al*, 2014; Price *et al*, 2014; Sousa-Victor *et al*, 2014; Tierney *et al*, 2014). Five thousand donor MuSCs were isolated from aged tamoxifen-treated Pax7Cre^ER^-tdTomato (tdT) mice, incubated for 120 minutes with either DMSO, ISR-i, or ISR-a and injected intramuscularly into a pre-injured wild-type adult mouse host. Recipient muscles were harvested 30 days after MuSC engraftment (Figure 6A). Quantification of tdT^+^ fibers showed that, compared to DMSO treatment, aged MuSCs treated with ISR-i lost transplantation potential, while treatment with ISR-a increased the output of transplanted MuSCs ∼2-fold (Figure 6B, C). Therefore, enforced ISR activity improved the physiology and *in vivo* regeneration function of aged MuSCs upon transplantation.

**Figure 6.**
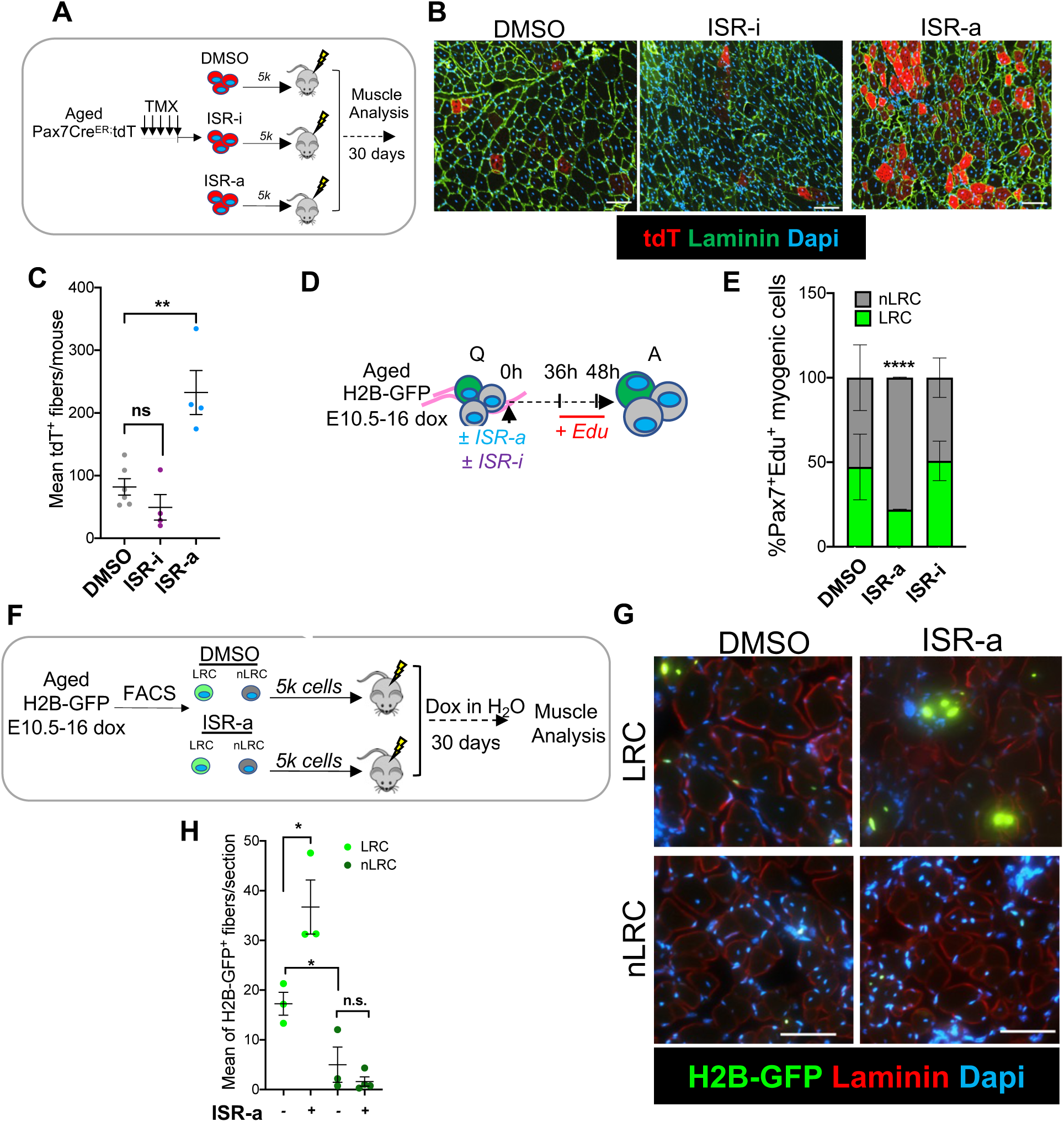
Sustained ISR is sufficient to rejuvenate aged MuSCs. **A.** Schematic of Pax7^ER/+^-tdT^+^ MuSCs, FACS isolated from a tamoxifen-treated aged Pax7Cre^ER^-tdTomato mouse, treated for 2 hours either with DMSO, ISR-a and ISR-i, and transplanted into a BaCl_2_-injured WT adult host. **B.** Representative images of muscle cross-section from transplanted Pax7^CE/+^-tdT^+^ pre-treated either with DMSO, ISR-a and ISR-i, stained for laminin (green) and DAPI (blue). Scale bar, 100µm. **C.** Mean of tdT^+^ donor-derived fibers/mouse from transplanted Pax7^ER/+^-tdT^+^ pre-treated either with DMSO, ISR-a and ISR-i. Mean±SEM; n=10->15 sections across 1mm of TA muscle, n=4-6 mice; **p<0.01. **D:** Schematic of single muscle fibers from embryonically doxed aged H2B-GFP mice (24 months) treated either with DMSO, ISR-a and ISR-i, pulsed with EdU and analyzed after 48 hours in culture. Q= quiescent, A= activated. **E.** Percentage of EdU^+^ myogenic LRC and nLRC at 48 hours after DMSO, ISR-a and ISR-i treatment. Mean±SEM; n=2 mice; ****p<0.0001. **F.** Schematic of donor LRC and nLRC MuSCs derived from embryonically dox-labeled aged (24 months) H2B-GFP mice, 2-hours treated upon FACS isolation with DMSO or ISR-a and transplanted into a BaCl_2_-muscle injured adult host. Dox water was given to the host during the 30 days of regeneration. **G.** Representative images of muscle cross-section from transplanted ±ISR-a LRC and nLRC, laminin (red) and DAPI (Blue). Arrows shows H2B-GFP^+^ centrally nucleated fibers. Scale bars, 100µm. **H.** Mean of H2B-GFP^+^ donor-derived fibers/mouse from transplanted ±ISR-a pre-treated LRC and nLRC from embryonically dox-labeled aged (24 months) H2B-GFP mice. Mean±SEM; n=10->15 sections across 1mm of TA muscle, n=4 mice; *p<0.05.

Both LRCs and nLRCs are capable of proliferation and commitment, however transplantation potential is limited to LRCs and aging is associated fewer LRCs and more nLRCs (Chakkalakal *et al*, 2012; Scaramozza *et al*, 2019). scRNA sequencing analysis confirmed that genes associated with ISR are enriched in nLRCs (Figure S6A). This is consistent with the overlap of ISR genes between transcriptional cluster 3 in total MuSC pool and nLRCs from adult muscle (Figure S1E).

To address the role of the ISR on MuSC subsets, we isolated single muscle fibers from aged TetO-H2B-GFP mice, that were embryonically labeled with doxycycline (Chakkalakal *et al*, 2012), cultured with ISR-a, ISR-i or DMSO, pulsed with EdU and fixed at 48 hours (Figure 6D). We found that ISR-a, but not ISR-i treatment, significantly increased the percentage of EdU^+^ MuSCs per muscle fiber compared to control, like adult MuSCs (Figure S6B; Figure 2A). ISR-a significantly enhanced the percentage of EdU^+^ cells within the nLRC fraction (Figure 6E) and increased overall EdU^+^ cells without affecting cell survival (Figure S6B, C). In contrast, ISR-i treatment decreased cell cycle entrance and increased apoptosis in both LRCs and nLRCs (Figure S6C). These data show that increasing ISR activity selectively accelerates activation of short-term MuSC progenitors (nLRCs).

Because transplantation potential is restricted to LRCs, we used a cell transplantation assay to test whether ISR activation enhances long-term stem cell capacity or reprograms short-term progenitors into LRCs. Five thousand LRCs and nLRCs were isolated from aged Tet-O-H2B-GFP mice, embryonically dox-labeled from E10.5 to E16. LRCs and nLRCs were incubated for 120 minutes with DMSO or ISR-a and injected intramuscularly into a pre-injured wild-type adult mouse host, under a dox-water regimen for 30 days to label engrafted cells (Figure 6F). In accordance with our previous work, quantification of H2B-GFP^+^ fibers showed that LRCs have increased stem cell capacity compared to nLRCs. Pre-treatment of aged MuSCs with ISR-a led to a 2-fold increase in the number of H2B-GFP+ fibers per section of LRCs, with no change in short-term progenitors (Figure 6G, H). Therefore, stimulation of the ISR pathway increased the output of aged long-term stem cells but did not reprogram short term progenitors in the transplantation model.

### Altering the ISR contributes to muscle regeneration *in vivo*

Given the enhanced myogenic capacity and transplantation potential associated with ISR activation, we sought to determine whether ISR activation could improve skeletal muscle repair, particularly in aged muscle. We administered ISR-a intraperitoneally to both adult and aged mice to transiently activate ISR, followed by barium chloride-induced injury to the Tibialis Anterior (TA) muscle, and mice were sacrificed at 21 days post-injury (Figure 7A). Both adult and aged mice in the ISR-a group exhibited a larger average cross-sectional area (CSA) and increased regenerating fiber sizes compared to aged vehicle-treated controls (Figure 7B). Notably, aged ISR-a-treated mice had even larger fibers than adult vehicle-treated mice (Figure 7C, D). Together, these results demonstrate that transient ISR activation enhances myofiber regeneration in both adult and aged mice.

**Figure 7.**
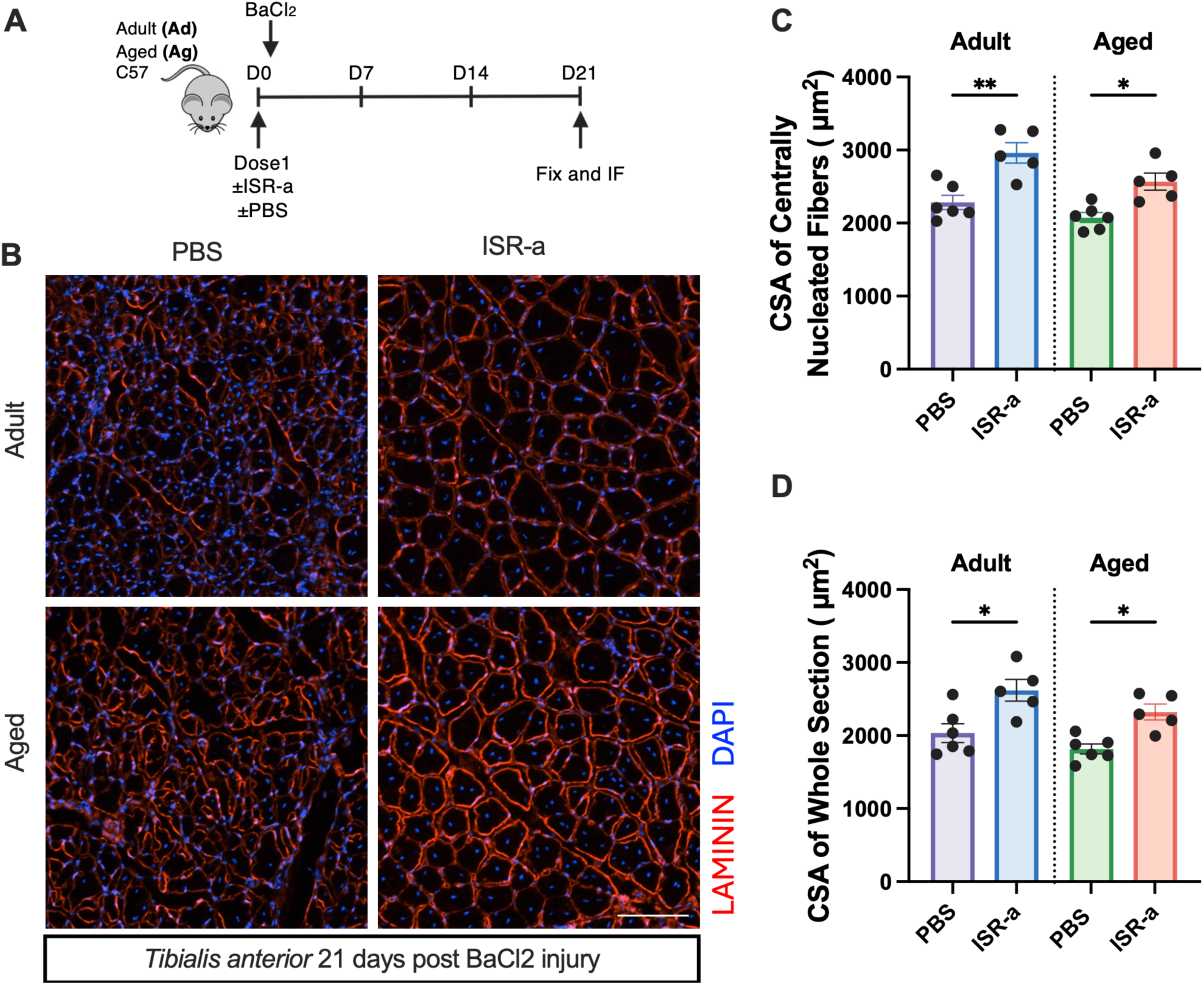
Altering the ISR contributes to adult and aged muscle regeneration *in vivo*. **A.** Schematic of IP injections of PBS and ISR-a followed by BaCl₂-induced injury into adult and aged mice TA muscle. TA’s were extracted at 21 days post-injury. **B.** Representative images of immunofluorescence analysis with antibodies against laminin (red) counterstained with DAPI (blue) on transverse sections of TA muscle from adult and aged C57 mice treated with PBS or ISR-a, 21 days after BaCl_2_-induced injury. **C.** Quantification of centrally nucleated myofiber cross-sectional area (CSA) in adult and aged mice with PBS or ISR-a. Mean±SEM; n=5/6. *p<0.05, **p<0.01. **D.** Quantification of myofiber CSA in the whole section in adult and aged mice with PBS or ISR-a. Mean±SEM; n=5/6. *p<0.05.

## DISCUSSION

Phenotypic and molecular analysis has revealed that MuSCs transit through a series of intermediate states as they exit quiescence, activate, and progress towards replication. Aged MuSCs follow the same activation trajectory, but at a slower rate. Therefore, the speed that MuSCs traverse through states is associated with its regenerative competence (Kimmel *et al*, 2020). We analyzed single cell transcriptional and behavior dynamics to identify regulators of MuSC function. This novel approach to study stem cell dynamics uncovered the role of the ISR pathway as a rejuvenator of aged MuSCs and muscle repair.

p-eIF2α is expressed in quiescent MuSCs, decreases during the first few hours of activation, and then reverses, surpassing levels found in the quiescent state. The variability of p-eIF2α levels within the MuSC pool increases during activation, suggesting a heterogenous activation response. These highly dynamic and heterogenous changes reveal a significant remodeling of the protein translation machinery during activation. Quiescent stem cells produce less protein than their immediate progenitors (Blanco *et al*, 2016; Signer *et al*, 2014, 2016; Zismanov *et al*, 2016). And, similar to hair follicle stem cells and HSCs (Blanco *et al*, 2016; Signer *et al*, 2014), our data show that the activation of MuSCs is not sufficient to reverse translational repression, based on the increased levels of p-eIF2α. We find that aged quiescent MuSCs have higher p-eIF2α levels compared to adult, suggesting an accumulation of stress during aging. Despite this, the ISR response to activation cues is blunted in aged MuSCs relative to adult, delaying MuSC activation. Treatment of MuSCs with an ISR activator increased p-eIF2α levels in adult and aged MuSCs to a similar degree, suggesting that p-eIF2α function is intact, but there may be age-dependent changes in the upstream sensing mechanisms.

During activation, MuSCs undergo metabolic changes, such as increased ATP, elevated mitochondrial activity and ROS, mTORC1 activation, enhanced autophagic flux, and induction of stress pathways including members of the ISR family, such as *ATF3* (Almada *et al*, 2021; L’honoré *et al*, 2018; Machado *et al*, 2017, 2021; Pallafacchina *et al*, 2010; Raval *et al*, 2020; Ryall *et al*, 2015; Tang & Rando, 2014; van den Brink *et al*, 2017; van Velthoven *et al*, 2017). The temporal relationship between these processes has not been resolved to date. Our gene ontology analysis revealed an enrichment of ISR-related genes in the early activated cluster 3 and genes associated with oxidative phosphorylation and ATP metabolic processes enriched in the later activated cluster 4, suggesting coupling between protein synthesis-based stress response and mitochondrial metabolism. ROS production is increased during MuSC activation (L’honoré *et al*, 2018) and may activate the ISR axis through the oxidation of ER proteins and activation of PERK (Harding *et al*, 2003; Zismanov *et al*, 2016). This coordination fine tunes protein translation when cells are exposed to oxidative stress. We also found that the early activated MuSC cluster 3 had increased levels of *Sqstm1*, also known as *p62*, an autophagic adapter to coordinate degradation of ubiquitinated substrates (Seibenhener *et al*, 2004). The coordination between autophagy and ISR provides the cell with an efficient mechanism to sense and clear misfolded proteins.

mTORC represents the other major pathway involved in the control of protein synthesis and survival upon nutrient availability, growth factor signaling, and stress (Laplante & Sabatini, 2012). Consistent with prior work, we here report that inhibition of general protein translation through rapamycin delayed MuSC activation (Zhang *et al*, 2015; Ge *et al*, 2009). Whereas ISR-a, which inhibits protein synthesis and increases selective translation through the transcription factor ATF4, accelerated adult and aged MuSC activation. Therefore, activation of the ISR is not dependent on general protein synthesis inhibition to promote a faster pace through activation states. ISR stimulates selective translation of proteins involved in amino acid import, oxidative stress, and the cell cycle (Fujita *et al*, 2021; Harding *et al*, 2003). However, further experiments are required to identify the downstream effectors of the ISR pathway that regulate the transition rate through MuSC activation state space.

Aged MuSCs experience dampened motility and cell behavior during activation which results in reduced regenerative capacity (Kimmel *et al*, 2020). We demonstrate that ISR activation in aged MuSCs partially restores many motile features involving speed, distance, and direction compared to adult MuSCs, specifically in the more motile cluster. These cell states are dictated by changes in single cell trajectories (Kimmel *et al*, 2020). Using cell painting, we further support these improvements in activation and behavior in ISR-a treated aged MuSCs by demonstrating higher DNA and mitochondrial content which concurs with later cell cycle phases and higher metabolic activity (Mapkar *et al*, 2025; Pala *et al*, 2018) and decreasing dysregulated non-homogenous actin structure, which is indicative of an aging state (Stearns-Reider *et al*, 2017). Our single cell sequencing results also indicated that activating the ISR in aged MuSCs drives a higher proportion of cells toward the more activated cluster, where more adult activated MuSCs reside, compromising of genes coinciding with mitochondrial metabolism and cell cycle. As MuSCs enter a more activated state, sustained energy is required during cell proliferation and activation (Sousa-Victor *et al*, 2022). Therefore, forcing higher ISR in aged MuSCs during activation could also lead to an increase in respiratory capacity.

MuSCs are heterogenous, with subsets transitioning through activation states more rapidly than others, such as Pax3^+^ MuSCs and LRCs (Kimmel *et al*, 2020). We found that ISR-a-mediated ISR activation impacted LRCs and nLRCs in a context-dependent manner. ISR-a increased activation rates of nLRCs *in vitro* with minimal effect on LRCs and enhanced the transplantation potential of LRCs with minimal effect on nLRCs. While there are technical differences between these assays, this data suggests that ISR stimulation increases activation speed in MuSCs with slower activation kinetics, such as aged MuSCs, but it does not reprogram short-term progenitors (nLRCs) into stem cells (LRCs).

Stem cell exhaustion and proteotoxic stress are two hallmarks of aging (López-Otín *et al*, 2013). Activation of the proteotoxic stress response represents a promising therapeutic approach to enhance stem cell function and increase resilience during aging. Our results revealed an increased proportion of regenerating fibers with larger fiber formation following ISR activation in aged mice, suggesting a prolonged regeneration window. A possible explanation is that the ISR-induced restoration of aged MuSC activation leads to a more efficient repair process. One potential mechanism underlying the difference in the regenerating window between ISR-activated adult and aged mice could be the age-related physiological repair environment. Aged muscle is typically more fibrotic and inflammatory, with an accumulation of extracellular matrix and increased pro-inflammatory cytokines (Moiseeva *et al*, 2023), whereas healthier adult muscle may already have an optimal regenerative capacity. Indeed, further investigation should consider if ISR-a could affect other muscle resident cell populations, such as fibroadipogenic progenitors and aged associated fibrosis.

In conclusion, our findings suggest that stimulating the ISR restores aged MuSCs to attain a more youthful activation dynamic, which improves regeneration. Importantly, we show that the ISR was required for adult MuSC activation, and this was blunted in aged conditions. By studying dynamical changes in molecular and phenotypic stem cell responses to activation cues we uncovered novel aspects of MuSC function and rejuvenation pathways. Future studies should endeavor to explain whether dynamically-regulated pathways play a role in other stem cell compartments during aging.

## ACKNOWLEDGEMENTS

We would like to thank the members of the Brack laboratory for critical discussions in the preparation of this manuscript. We acknowledge Xuefeng Sun for technical assistance. We acknowledge the Parnassus Flow Cytometry Core and Genomics CoLab Core at UCSF, supported in part by Grant NIH P30 DK063720 and NIH S10 1S10OD021822-01. This work was supported by NIH grants (R01AR060868, R21AG063416) to ASB and Calico-QB3 Longevity Fellowship to AS.

## AUTHOR CONTRIBUTION

ADB and AS conceived the project, designed and performed experiments, analyzed data, interpreted results and wrote the manuscript. HZ performed *in vivo* injury experiments. TT analyzed sc-RNA sequencing data and interpreted results. SM analyzed sc-RNA sequencing and performed cell painting experiments. NS analyzed heteromotility data. SE performed experiments and analyzed results. BF and XL interpreted experiments and edited the manuscript. ASB conceived the project, interpreted experiments and wrote the manuscript.

## Data and Code Availability

Raw single cell data is available in GEO under accession number GSE143476 and GSE306935. Code availability for:

Heteromotility: https://github.com/cellgeometry/heteromotility

Cell painting: https://github.com/samihamahin/ISR_cellpainting

scRNA sequencing: https://github.com/samihamahin/ISR-singlecellRNAsequencing

## METHODS AND MATERIALS

### Animals

Animals were handled according to UCSF Institutional Animal Care and Use (IACUC) guidelines. Experimental mice were housed in a pathogen-free barrier facility. Mice were also housed under a 12-hour light/12-hour dark cycle and temperature-controlled environment with standard diet and water ad libitum. Adult (2-5 months) and aged (18-24 months) C57Bl/6J (C57) were purchased from Jackson Laboratory. TetO-H2B-GFP were embryonically labeled by administration of doxycycline from E10.5-E16 and chased until aged (18-24 months). Pax7Cre^ER^tdT (18-24 months) were injected with tamoxifen for five days. Tamoxifen was resuspended in coin oil at 20 mg/ml and administered at 150 mg/kg body weight via intraperitoneal (IP) injection. All animals were maintained on a C57 background.

### Single cell RNA-Sequencing

We used previous datasets from our laboratory (Kimmel *et al*, 2020) where adult (3-5 month) and aged (20-24 month) C57 or TetO-H2B-GFP MuSCs were isolated by FACS. MuSCs were either sequenced straight away (quiescent) or after 18 hours of activation (activated). In a separate dataset, adult (4 months) and aged (22 months) MuSCs were isolated by FACS and plated on extracellular coated plates (ECM) in growth media ((F10 (Gibco), 20% fetal bovine serum (FBS) (Gibco), (5 ng/ml) FGF2 (R&D Systems)) for 18 hours in DMSO or 10 µM Sal003 (activated). The activated cells were dissociated using cell dissociation buffer (Gibco), stained with propidium iodide and FACS sorted to remove dead cells and debris. Both live quiescent and activated cells were transferred to the 10x Chromium System for library preparation and sequencing using Illumina NovaSeq.

### Single cell RNA-sequencing analysis and pseudotime analysis

The single-cell RNA-sequencing analysis was described previously (Kimmel *et al*, 2020). Subsequent fastq files were processed using the count function from CellRanger (version 5.0.1) and data analysis was carried out using the Seurat package (version 3.1.1) in R (version 3.6.1) (Butler *et al*, 2018; Stuart *et al*, 2019). Suspected dead or doublet cells were filtered out using the following criteria: >10% mitochondrial genes and >5,000 expressed genes/cell. Transcripts Gm42418 and AY036118, which overlap an unannotated Rn45s rRNA locus, were removed before normalization. Gene expression was normalized by the “NormalizeData” function with a scale factor 10,000 and log-transforming. Top 2,000 variant genes in the dataset were identified using the “FindvariableFeatures” function with the vst method. After the scaling of the expression of each gene, we performed PCA on the variable genes and used the first 10 PCs for subsequent clustering and visualization. Unsupervised clustering was performed using a graph-based approach implemented in the Seurat with low resolution (0.2). Non-linear dimensional reduction by UMAP was performed using the first 10 PCs to visualize the dataset. We removed four small clusters that we deemed to be non-myogenic cells based on the low expression of *PAX7* and *MYOD1*. To identify differentially expressed (DE) genes, the non-parametric Wilcoxon rank sum test was performed using the “FindMarkers” function in Seurat. Genes with adjusted p-value <0.05 and |logFC| >0.25 were considered as differentially expressed. Gene ontology enrichment analysis was performed with DAVID GO (Huang *et al*, 2009; Sherman *et al*, 2022) and considered terms for biological processes and KEGG Pathways. Psuedotime analysis was performed with the Monocle3 package (Cao *et al*, 2019). The preprocessed Seurat object was converted into a cell data set using the method “as.cell_data_set”. And the pseudotime graph was generated using the methods “order_cells” and “plot_cells”.

### MuSCs isolation by FACS

Skeletal muscle tissue from mouse hindlimb and forelimb was dissected and digested with 0.2% collagenase II at 37°C for 90 minutes. Upon washing, the muscle was digested in 0.2% collagenase II and dispase 0.4% for 30 minutes at 37°C. Digested tissue was passed through a 20-gauge needle, serially filtered through a 40 and 20 µm filter and centrifuged at 1500 rpm at 4°C for 5 minutes, to obtain a single cell suspension. Muscle stem cells were stained and FACS sorted using a triple negative CD31/CD45/Sca1 (Ly-6A/E) and a double positive V-CAM (CD106)/integrin-α7 selection. Flow cytometric sorting was performed on a FACS-Aria II (BD Biosciences).

### Single muscle fibers isolation and culture

Single muscle fibers were isolated from EDL (Extensor Digitorum Longus) muscle of adult (3-5 months) and aged (18-24 months) mice upon 0.2% collagenase II dissociation for 1.5 hours at 37°C. Dissociated single muscle fibers were manually collected and cleaned under a dissection microscope, then either fixed in suspension immediately in 4% PFA for 10 minutes or cultured for 3, 6, 18, 24, 48 and 72 hours with plating media (Dulbecco’s Modified Eagle’s Medium (DMEM) and 10% horse serum (HS)). Sal003 and ISRIB were dissolved in DMSO and used respectively at 10 µM and 200 nM final concentration in plating media. For activation experiments, EdU (Carbosynth) was pulsed into the media at 12 and 2 hours (10 μM) prior to fixation with 4% PFA for 10 minutes. Muscle fibers were then processed for EdU staining and immunofluorescence.

### Primary mouse myogenic cells *in vitro* culture

For EdU and survival experiments FACS MuSCs were plated either on 8-well chamber slides or 96 well plates pre-coated with ECM and cultured with growth media for 48 hours. Prior to fixation at 48 hours, EdU was pulsed into media at 12 and 2 hours. For survival, after 48 hours MuSCs were fixed and stained for Caspase 3. FACS-purified MuSCs were cultured in vitro either with DMSO, Sal003 at 10 µM, ISRIB at 200 nM, Rapamycin at 10 µM and CHX at 100 µg/ml (Zismanov *et al*, 2016) in high serum media for 3, 24 or 48 hours. To inhibit general protein synthesis FACS-purified satellite cells were cultured *in vitro* either with DMSO, Rapamycin or CHX in growth media for 3 hours. For CHX experiment, we then washed out the cells for 45 hours. Cells were then fixed and processed for EdU staining and immunofluorescence.

### EdU Staining and Immunohistochemistry

Single fibers and FACS sorted MuSCs were fixed with 4% PFA for 10 minutes and stained using the Click-iT EdU Alexa Fluor 647 kit protocol (Thermo Fisher Scientific). EdU was detected by fluorescence microscopy. For immunohistochemistry, cells/fibers were fixed with 4% paraformaldeyde for 10 minutes at room temperature, washed with PBS three times for 5 minutes, permeabilized for 10 minutes with 0.2 % Triton-100x (Sigma-Aldrich) and then blocked with 10% goat serum in PBS-TX for 30 minutes. Cells were then incubated with the specific primary antibody in blocking solution overnight at 4°C. Corresponding species-specific secondary antibody and 100 ng/ml DAPI (4,6 diamino-2-phenylindole; Vector Laboratories) were applied for 1 hour at room temperature in blocking solution. Fibers were finally washed and mounted with Fluoromount (Southern Biotech). Images were processed with Nikon NID-elements software.

### Lentiviral Infection

FACS-purified MuSCs were plated at 2500 cells/well in growth media, treated with DMSO, Sal003 and after 18 hours infected with lentivirus containing GFP, scrambled control or shRNA to *Atf4* and *Atf4* shRNA with Sal003 according to manufactures protocol (Vector Builder) at 50 Multiplicity of Infection (MOI) for 24 hours. Cells were then rinsed and incubated in fresh growth media for 6 hours, fixed and processed for EdU staining and immunofluorescence.

### Time-lapse Imaging and cell behavior analysis

MuSCs were plated (2500 cells/well) and imaged on a 96 well plate on an incubated microscopy platform (Oko lab) for 48 hours. DIC images were collected every 6.5 minutes and used for tracking cellular motility and behavior with *Heteromotility* as described previously (Kimmel *et al*, 2018).

First, we identify and segment cells using a deep learning-based instance segmentation algorithm that returns the coordinates and outlines of cells. The segmentation algorithm is typically trained to recognize objects on smaller images. However, our well images are much larger and have a high resolution. To use the model effectively while preserving resolution, we divide each well into 6×6 grids. Each grid is then fed into our segmentation pipeline to identify and segment cells. We use a detectron2-ResNeSt-based model pre-trained on the LIVECell dataset (Edlund *et al*, 2021) for instance segmentation. This model saves the (x,y) coordinates for every cell in each grid of the well across all the video frames. We then stitch the grids and extracted (x,y) coordinates for each cell back together for the entire well before the tracking step. For tracking, we use a SORT-based multi-object tracker (Kimmel *et al*, 2018). This algorithm uses the past locations of cells to estimate their current location using Kalman filters and matches cells across frames using the Hungarian matching algorithm. The results are saved as (x,y) coordinates of tracked objects across video frames. We use these coordinates to extract motility-based features (Kimmel *et al*, 2018).

### Cell Painting Assay and Quantification

MuSCs were plated on ECM coated 384 well plates with Sal003 or DMSO and cultured for 18 hours in growth media and fixed. Cells were stained using PhenoVue™ Cell Painting Kit (Revvity) in line with previous protocols (Cimini *et al*, 2023; Bray *et al*, 2016) with small modifications to visualize the mitochondria, f-actin cytoskeleton and nucleus. Thirty minutes prior to fixation, PhenoVue 641 mitochondrial stain (272 μg/ml) was added to live cells. Post fixation cells were washed and permeabilized in 0.2% triton X-100 with PBS for 20 minutes. Cells were washed in PBS and PhenoVue 568 phalloidin (48 μg/ml) and Hoechst 33342 (5 μg/ml) dyes were incubated in 1x PhenoVue Dye Diluent A and 0.2% (vol/vol) triton X-100 for 30 minutes at room temperature. Cells were washed and stored at 4°C until imaging. Confocal imaging at 63x magnification was performed on the Opera Phenix High Content Screening system (Revvity) and data are generated with the Harmony software (Revvity).

The data was normalized with pyctominer’s (Serrano *et al*, 2025) standardize function and scaled with scikit-learn’s RobustScaler. Features were selected by scikit-learn’s the univariate selection method SelectKBest and scored using ANOVA’s F-test to find significant features varying across groups and the means for each feature are plotted along with the F-value. The cells were clustered using unsupervised Gaussian Mixture Models (GMM) and Akaike information criterion (AIC) and Bayesian information criterion (BIC) were used to determine the number of clusters for the GMM model. PCA was performed and clusters were trained using a Random Forrest Classifier (Breiman, 2001) to determine the most important features for each cluster and generated feature importance with the mean decrease impurity (MDI).

### MuSC transplantation

For transplantation experiments, MuSCs from Pax7Cre^ER^tdT mice (tdT^+^Vcam^+^/Int-a7^+^/CD45^-^CD31^-^/Sca1) and LRC and nLRCs from embryonically doxed *TetO-H2B-GFP* mice were pre-treated with DMSO, Sal003 or ISRIB during FACS sorting (2 hours) and injected into adult TA muscle (5000 cells/mouse) of C57 mice that had been injured 1.5-days before with 1.2% BaCl_2_ (Sigma).

### *In vivo* Drug Administration and Injury

Adult (4 month) and aged (20-22 month) C57 mice were administered 1-hour pre-BaCl_2_ injury via IP either Sal003 (HY-15969, MedChemExpress, 1 mg/kg) or ISRIB (HY-12459, MedChemExpress, 1 mg/kg), both dissolved in DMSO and PBS. Twenty-one days after injury, the mice were euthanized and TA muscles were extracted, frozen in 2-methyl isobutane over dry ice, and stored at -80 for histology and immunofluorescence.

### Histology and immunofluorescence

TA muscles were cryo-sectioned (8 µM thickness) onto glass slides and sections underwent blocking with 10% Goat Serum, 0.2 % PBS-TX and then incubated with rabbit anti-laminin (1/500, Abcam) overnight at 4°C. Anti-rabbit secondary antibody and DAPI were applied for 1 hour at room temperature, then washed and mounted for microscopy analysis.

### Quantification and Statistical Analysis

All results are presented as mean ± standard error of the mean (SEM) or standard deviation (SD) where indicated. Statistical analysis was performed using a student’s t-test to calculate differences between two groups or simple linear regression to calculate the correlation between two variables. Analysis was conducted using one-way or two-way ANOVA with Tukey’s post-hoc correction for multiple comparisons. Sample size and/or replicate number for each experiment are indicated in the figure legends. Results with p-values of less than 0.05 were considered statistically significant. All statistical analysis was quantified using Graphpad Prism® (version 10).

