## Supplementary File 1 for "The Integrated Stress Response Pathway Improves Aged Murine Muscle Stem Cell Activation and *in vivo* Regeneration"

### Supplementary Figures

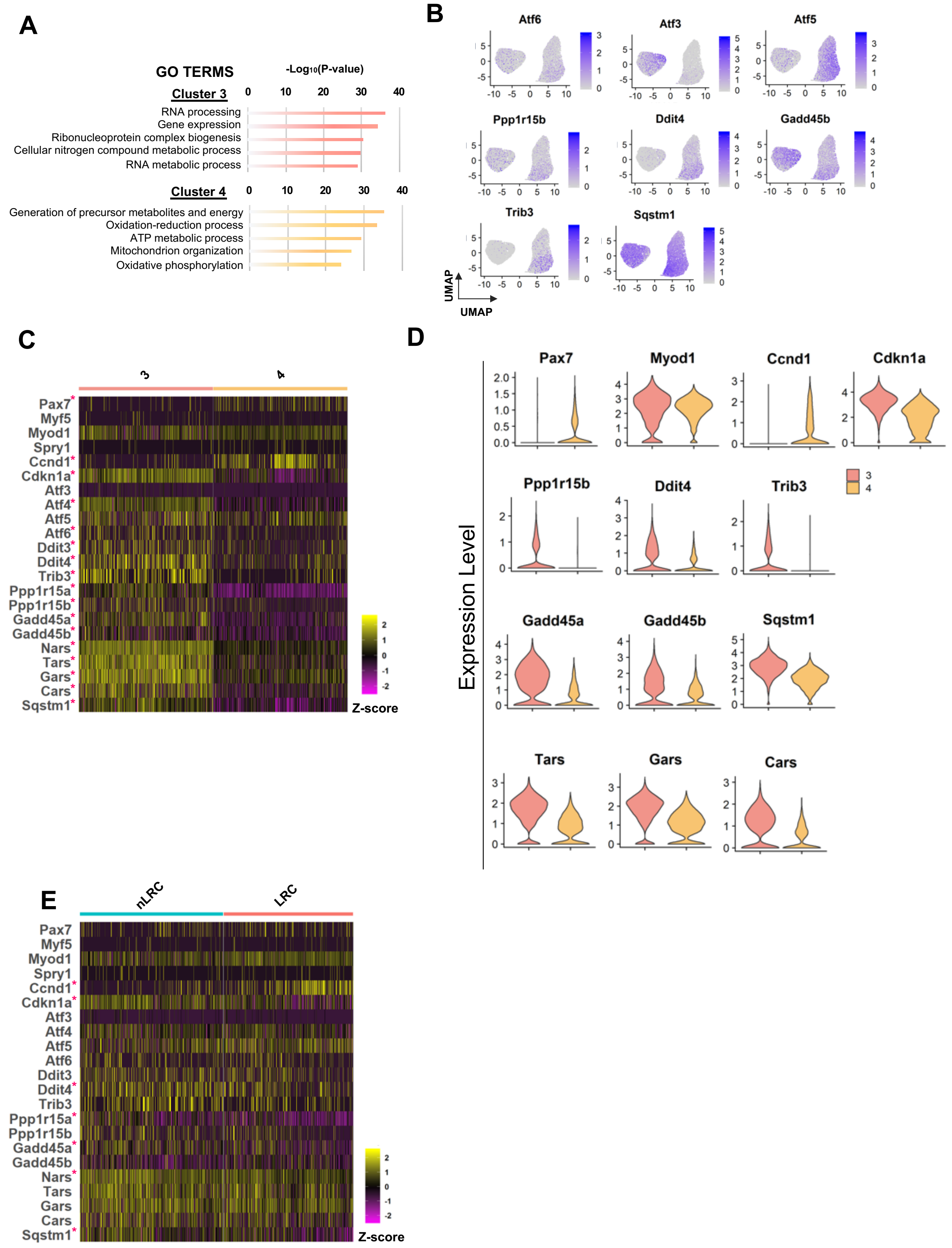

**Figure S1. The ISR pathway undergoes dynamic expression changes during adult MuSC activation.**

**A.** Gene ontology analysis for common genes with statistically significant differentially expressed genes among cluster 3 (top) and cluster 4 (bottom) was performed using g:Profiler, only considering biological processes.
**B.** Overlay of expression of ISR-related genes in scaled log (UMI+1) on the UMAP plot.
**C.** Heatmap of myogenic, cell cycle and ISR pathway genes in cluster 3 and cluster 4 of activated adult MuSCs. Asterisks indicate statistically significant ($p<0.05$) differences between clusters. The color in the heatmap represents Z-score normalized expression for each gene.
**D.** Violin plots depicting differential myogenic, cell cycle and ISR-gene expression levels between cluster 3 and cluster 4.
**E:** Heatmap of myogenic, cell cycle and ISR pathway genes in nLRCs and LRCs of activated adult MuSCs. Asterisks indicate statistically significant ($p<0.05$) differences between clusters. The color in the heatmap represents Z-score normalized expression for each gene.

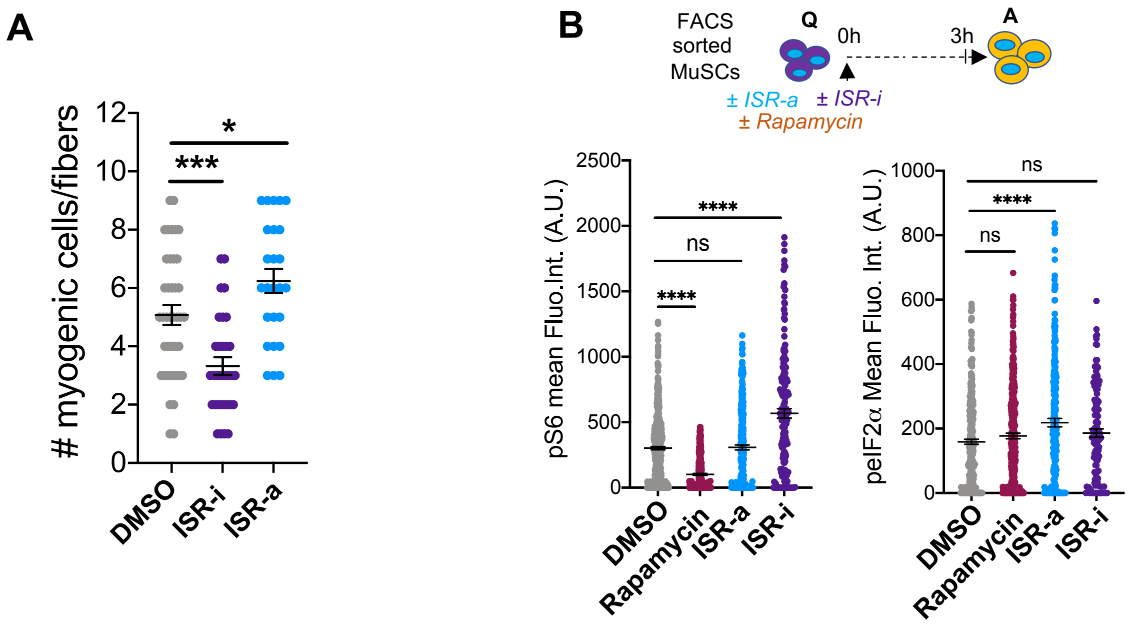


**Figure S2. Stimulating the ISR pathway is associated with accelerated adult MuSC activation through selective translation.**

**A.** Number of myogenic cells per muscle fibers after 72 hours culture with treatment. Mean±SEM; n=2; *p<0.05; ***p<0.001.

**B.** Schematic of FACS sorted MuSCs *in vitro* cultured in DMSO, rapamycin, ISR-a and ISR-i for 3 hours (top). pS6 (bottom left) mean and peIF2α (bottom left) fluorescence intensities in FACS sorted MuSCs cultured for 3 hours after isolation in the presence either of DMSO, Rapamycin, ISR-a and ISR-i. Mean±SEM; n=2 mice; ****p<0.001, ns=not significant.

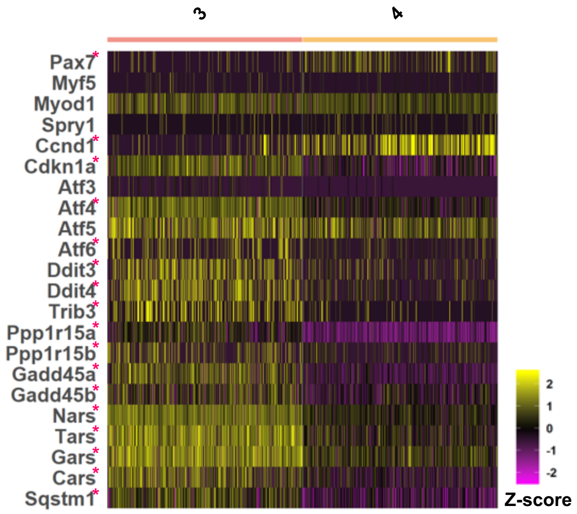


**Figure S3. Activation of ISR is sufficient to restore function of aged MuSCs.**

**A.** Heatmap of myogenic, cell cycle and ISR pathway genes in cluster 3 and cluster 4 of aged activated MuSCs. Asterisks indicate statistically significant ($p<0.05$) differences between clusters. The color in the heatmap represents Z-score normalized expression for each gene.

**A**

| Feature Name | Notation | Description |
|---|---|---|
| Total Distance | Total Distance | Total distance traveled |
| Net Distance | Net Distance | Net distance traveled |
| Maximum Speed | Max Speed | Maximum speed |
| Average Speed | Avg Speed | Average overall speed |
| Time Spent Moving | Time Moving | Proportion of time in motion, variable time intervals considered [1,10] frames |
| Average Moving Speed | Avg Moving Speed | Average speed during movement, variable time intervals considered [1,10] frames |
| Linearity | Linearity | Pearson's $r^2$ of regression through all cell positions |
| Progressivity | Progressivity | (net distance traveled / total distance traveled) |
| Hurst Exponent | Hurst RS | Hurst exponent estimation using Mandelbrot's rescaled range methods |
| Autocorrelation | Autocorr | Autocorrelation of the displacement distribution for variable time lags |
| Proportion of Right Turns | P Turn | Proportion of turns an object makes to the right, calculated for multiple time lags and direction estimation intervals |
| Maximum Turn Magnitude | Max Theta | Maximum turn angle in radians, for various time lags and direction estimation intervals |
| Average Turn Magnitude | Mean Theta | Mean turn angle in radians, for various time lags and direction estimation intervals |
| Random Walk Net Distance | rw_netdist | (cell path net distance - simulated random walk net distance) |

**B**

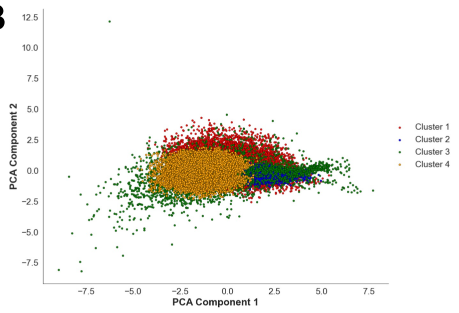


**C**

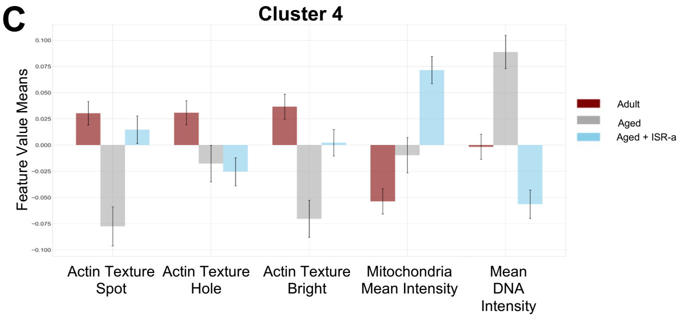


**D**

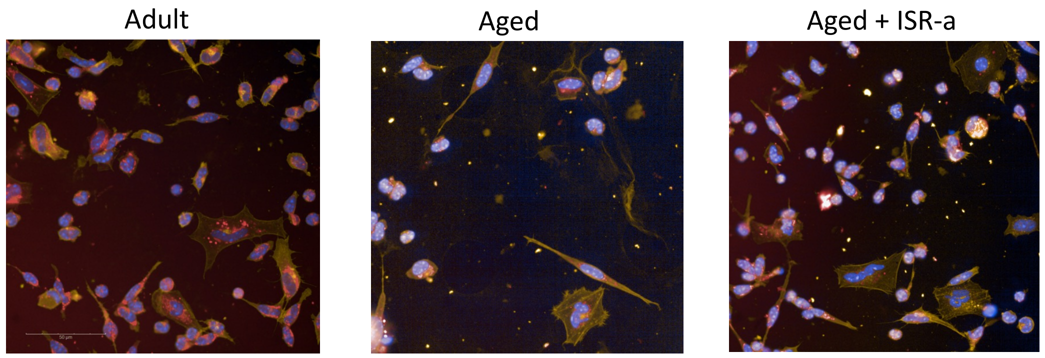

**Figure S4. Aged MuSC behavior was improved with ISR activation through alterations in actin and mitochondria morphology.**

**A:** Table of feature names, notations and feature descriptions.
**B.** PCA projection of morphological state space and unsupervised GMM clustering identifies four unique clusters.
**C.** Histograms of the most variable features means in cluster 4 between adult DMSO, aged DMSO and aged ISR-a
**D.** Representative confocal images from the cell painting assay in adult DMSO, aged DMSO and aged ISR-a. Overlapping mitochondrial (red), actin (yellow), and DNA (blue) stains. Scale = 50 µm. n=3.

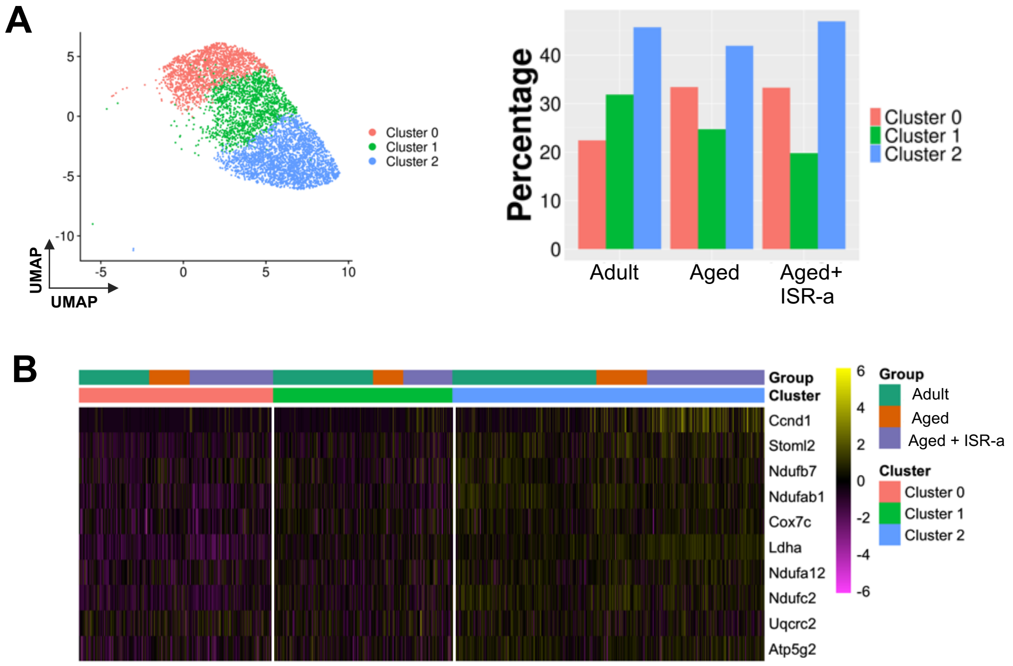


**Figure S5. Adult and aged activated MuSCs show differences of activation correlated with ISR activity.**

**A.** UMAP projection of unsupervised clustering showing 3 transcriptional clusters within activated adult, aged and aged with ISR-a cell populations (left). Percentage of clusters with adult, aged and aged with ISR-a (right).
**B.** Heatmap of upregulated metabolism genes in adult, aged and aged with ISR-a groups and the 3 transcriptional clusters.

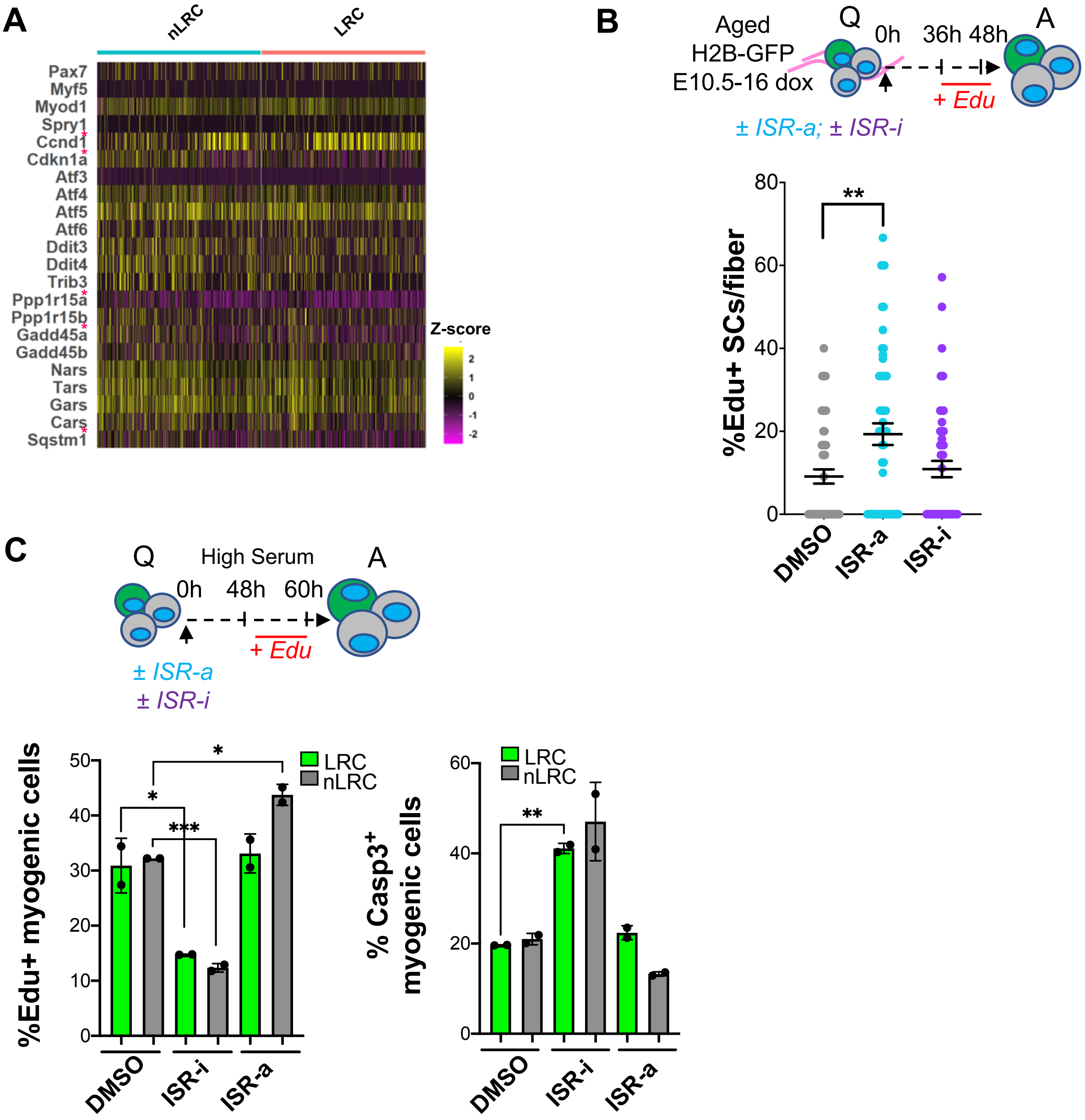


**Figure 6. The ISR differentially regulates subsets of the aged MuSC pool in a context dependent manner**

**A.** Heatmap of myogenic, cell cycle and ISR pathway in nLRC and LRC of activated aged MuSCs. Asterisks indicate statistically significant ($p<0.05$) differences between the groups. The color in the heatmap represents Z-score normalized expression for each gene.

**B.** Schematic of single muscle fibers from embryonically labeled aged H2B-GFP mice, cultured in the presence or absence of ISR-a and ISR-i, pulsed with Edu at 36 hours and analyzed at 48 hours (top). Percentage of Edu$^{+}$ MuSCs per muscle fibers after 48 hours in culture with either DMSO, ISR-a and ISR-I (bottom). Mean±SEM; n=2; **p<0.001

**B.** Schematic of FACS sorted MuSCs *in vitro* cultured in presence or absence of ISR-a and ISR-i, pulsed with Edu at 48 hours and analyzed at 60 hours (top). Percentage of Edu$^{+}$ (bottom left) and Casp3$^{+}$ (bottom right) myogenic cells after 60 hours in culture with DMSO, ISR-a and ISR-i. Mean±SEM; n=2; *p<0.05, **p<0.01, ***p<0.001.

#### Supplementary Methods and Materials

##### FACS antibody

For FACS cell isolation we used the following antibodies: anti-CD31 (Clone 390, PE-Cy7, 1:500, BD Pharmingen, 561410 AB10612003); anti-CD45 (Clone 30-F11, PE-Cy7, 1:500, BD Pharmingen, 552848 AB394489); anti-Ly6A/E (Clone D7, APC-Cy7, 1:500, BD Pharmingen, 560654 AB1727552); anti-Vcam1 (Clone 429, PE, 1:100, Invitrogen, RMCD10604 AB255657); anti-α7-integrin (Clone R2F2, APC, 1:500, Ab Labs, 67-0010-05).

##### Immunohistochemistry antibody

For immunohistochemistry staining we used the following antibodies: mouse anti-Pax7 (1/100, DSHB), mouse anti-cyclin D1 (1/250, Abcam), rabbit anti-peIF2α (1/100, Abcam), rabbit anti-Myod (1/150, Santa Cruz), rabbit anti-Myogenin (1/100, Santa Cruz), rabbit anti-Caspase 3 (1/500, Cell Signaling) and rabbit anti-pS6 (1/300, Cell Signaling). For secondary antibody we used: Alexa Fluor-conjugated 488, 546 and 647 at 1/2000 (Thermo Fisher).

##### Analysis of satellite cells and muscle fibers *in vivo*

The total number of Pax7$^+$tdTomato$^+$ and H2B-GFP$^+$ muscle fibers was quantified in a minimum of five to ten serial sections per muscle in three separate regions from the mid-belly of the muscle.

##### Image Processing and Quantification

All images were processed and edited using Nikon NIS Elements and ImageJ, applying modifications uniformly across the entire image. Myofiber cross-sectional area was determined using a CNN-based deep learning segmentation model for injury-specific myofiber analysis in Cellpose, with manual adjustments on immunofluorescent laminin-stained frozen sections of the TA. For each mouse, the cross-sectional areas from at least two sections were quantified from 10x magnification.
